# Modelling genotypes in their microenvironment to predict single- and multi-cellular behaviour

**DOI:** 10.1101/360446

**Authors:** Dimitrios Voukantsis, Kenneth Kahn, Martin Hadley, Rowan Wilson, Francesca M. Buffa

## Abstract

A cell’s phenotype is the set of observable characteristics resulting from the interaction of the genotype with the surrounding environment, determining cell behaviour. Deciphering genotype-phenotype relationships has been crucial to understand normal and disease biology. Analysis of molecular pathways has provided an invaluable tool to such understanding; however, it does typically not consider the physical microenvironment, which is a key determinant of phenotype.

In this study, we present a novel modelling framework that enables to study the link between genotype, signalling networks and cell behaviour in a 3D microenvironment. To achieve this we bring together Agent Based Modelling, a powerful computational modelling technique, and gene networks. This combination allows biological hypotheses to be tested in a controlled stepwise fashion, and it lends itself naturally to model a heterogeneous population of cells acting and evolving in a dynamic microenvironment, which is needed to predict the evolution of complex multi-cellular dynamics. Importantly, this enables modelling co-occurring intrinsic perturbations, such as mutations, and extrinsic perturbations, such as nutrients availability, and their interactions.

Using cancer as a model system, we illustrate the how this framework delivers a unique opportunity to identify determinants of single-cell behaviour, while uncovering emerging properties of multi-cellular growth.

**Availability and Implementation:** Freely available on the web at http://www.microc.org. Research Resource Identification Initiative ID (https://scicrunch.org/): SCR 016672

## Introduction

A comprehensive understanding of living organisms, including their development and the occurrence and progression of disease, requires systematic efforts into deciphering the link between the multitude of different genotypes and phenotypes, that is the set of observable characteristics resulting from the interaction of the genotype with the surrounding environment, determining cell morphology and behaviour.

Efforts to characterise such relationships have proliferated in recent years, thanks in part to the increased capability to efficiently collect the necessary data in an ever higher number of organisms and individuals. Such efforts have spanned from cataloguing genetic variation in thousands of individuals [see e.g. (1)] and searching for genotypes-phenotypes associations, to silencing or inactivating thousands of genes in laboratory high-throughput screens to study their function [see e.g. (2)]. However, efficient instruments to achieve a comprehensive understanding of the causal nexus between a given genotype and the observed phenotype are still lacking, and this is particularly true when such a nexus is complex and multifactorial (3-5). One promising means to achieve such an understanding has been the characterisation and modelling of biological pathways (5).

Several of the key biological pathways regulating cellular function are increasingly understood, along with their dysregulation in multiple diseases (6-9). However, much less is known about how these pathways interact and determine the behaviour of individual cells and multi-cellular systems. This has fuelled methodological development of efficient representations of such interactions in order to facilitate the study of the underlying potential mechanisms. A number of different approaches have been proposed, including modelling by differential equations, and network modelling methods, such as petri nets and logical networks (10-12). Such methods have produced encouraging results (13-16), but it is becoming increasingly evident that modelling molecular pathways and signalling, or gene, networks in isolation, dissociated from the cellular context, does not reflect the crucial impact of the microenvironment in determining the phenotype (17-19). Additionally, as single-cell sequencing and imaging technologies are providing new in-depth information about the genotype and phenotype of single, or small groups of, cells (20), modelling approaches that enable consideration of cells both as independent entities and as a population, are becoming increasingly attractive.

To address the above needs, we have developed a novel computational framework which combines Agent-Based Modelling (ABM) and gene network modelling. This framework tackles the fundamental challenge of integrating genotype with phenotype data, while accounting for certain important aspects of the physical microenvironment, a key determinant of the phenotype. The most innovative aspect of this framework is that it enables to build models of the genotype-phenotype relationship in a 3D spatially-aware microenvironment, including aspects such as signalling to and from the microenvironment, and signalling between cells. Importantly, the collective behaviour and evolution of cellular populations emerges from the properties and behaviour of individual cells, which in turn are governed by the underling dynamics of specific signalling networks, and the interactions with the surrounding cells and microenvironment. This permits to predict the behaviour of individual cells, and the entire population of cells, and it also enables to study of the possible causative mechanisms of such behaviour.

In this paper, we introduce this modelling framework, illustrate the capabilities of microC, a first cloud-based implementation of the framework, and we illustrate the range of potential applications that it enables. Specifically, we perform perturbation experiments of increasing complexity, in which we monitor over time the three-dimensional (3D) growth and evolution of mixed populations of cells.

To achieve this, we chose the example of cancer: a complex disease where methods to study the link between genotype and phenotype are particularly and urgently needed (4). To inform our choices for the gene network and the model parameters, we exploited previously acquired data on gene interactions and cell growth from a number of independent publications. We then built our initial model in a mechanistic “bottom-up” fashion, progressing from the individual elements to the whole system. Following this strategy, we simulated the 3D growth of cell spheroids focusing on pathways underlying the main hallmarks of cancer, including sustained proliferative signals, resistance to cell death and evasion of growth suppressors (9).

We thus considered a set of alterations amongst the most frequently observed across all cancer types, namely EGFR activation, p53 loss-of-function and PTEN loss-of-function (21). We then asked to what extent this initial model, which has been built based on general assumptions and not optimized for a specific cancer type, could reproduce experimental results not used for the model construction and obtained in a cancer which often displays these mutations. For this we chose basal-like triple negative breast cancer. This type of breast cancer lacks receptors for the hormones oestrogen and progesterone, and the Her2 protein, and thus is not responsive to treatments targeting these. Importantly, an immortal though not tumorigenic mammary tumour cell line exists, MCF10A, which is considered a suitable pre-malignant model (22), allowing us to assess the effect of inducing these alterations as single drivers, or together.

With this model, we gradually increased the complexity of our simulations to investigate the effect that varying genetic and microenvironmental parameters has on the model predictions, and on the resulting clonal competition, signalling to and from the microenvironment, and cell-cell interaction.

## Methods: the microC framework

### Rationale for the framework

Previous studies have modelled cells as computational agents and started to consider replacing ABM conditional statements, that drive cellular behaviour, with gene networks, that can represent more realistically the dynamical features of the intra-cellular system. In particular, small scale predefined logical networks have been used to model the cell-cycle arrest in a model of avascular spheroid growth (23), and to study cell differentiation in a hyper-sensitivity reaction model (24). However, these networks were considered as static entities, providing simple rules and not fully embedded in the ABM simulation. In another example, cells have been equipped with relatively more complex decision making models, represented as differential equation (25), but this approach was also not fully embedded in the ABM modelling, and is further limited by the requirement of prior knowledge for the many kinetic parameters to represent pathways accurately.

microC fully exploits ABM technology. Specifically, the cellular behaviour is determined by a gene network, encapsulated within the cell, and by the interactions of the gene network with the surrounding microenvironment (Figure 1). Both the cells and the inner networks are simulated using ABM. As a result, cellular behaviour may be studied both at the single-cell level, decoding the link between a cell genotype and its phenotypic realization in a given contextual environment, but also at the more global level, allowing the search for emerging properties of the multi-cellular system.

**Figure 1.**
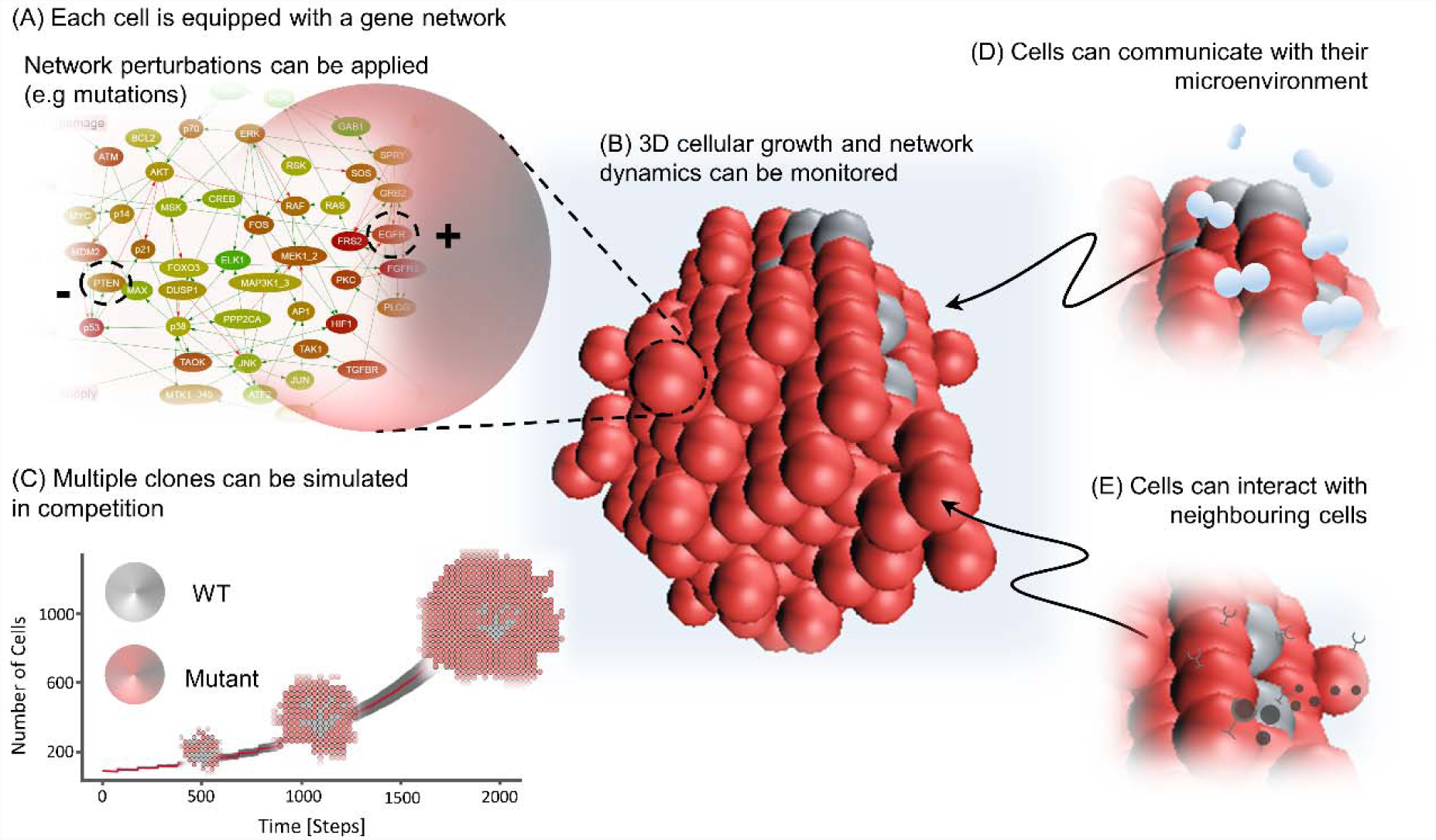
Conceptual representation of microC environment and simulation components. The microC framework enables multiscale simulations linking genotype to phenotype, via gene networks, by fully exploiting agent-based modelling. The technical details on the framework and its implementation are provided in the Methods. Here, we illustrate a study where four setups of increasing complexity are considered, evaluating the impact of new elements in a controlled fashion. A. We simulate the effect of introducing gene network perturbations (activating, +, or inactivating, -, mutations) in single clones B. We grow mono-clonal or multi-clonal populations of cells in a 3D environment (spheres visually represent single cells) C. We study competition between different clones grown together (colours represent different mutation profiles; the growth curve demonstrates exponential growth of aggressive clones; 2D simulation show growth patterns under competition) D. We enable interaction with the surrounding microenvironment, e.g. oxygen concentration variation that results to necrotic cells E. We enable signalling between cells (represented here as tiny black spheres produced by activated cells; junctions represent receptors).

### Key elements of the framework

To define a specific model in the microC modelling framework, a number of inputs and parameter values need to be provided. These define the environment of the simulation, some of the cell’s characteristics, the number of replicates considered in each simulation and the length of the simulation, the choice of the gene network, the cell mutation profiles, and how a cell interacts with the environment and other cells. These parameters, and how they define the specific models, are presented below and further discussed in the Supplementary Protocol and online documentation (http://www.microc.org). Of note, in this work, we set the parameter values for the individual elements of the model based on previously published data (see sections below). We then carry out sensitivity analysis to understand the implications of these choices for our specific case.

### Cells as computational agents

Cells are represented in microC as computational agents acting and interacting in a three-dimensional (3D) space. Cells are arranged in a rectangular 3D grid where each voxel may be occupied by only one cell. They may interact with the microenvironment by consuming oxygen, and with other cells, via diffusible substances, such as cytokines, chemokines and growth factors/hormones, whose production is defined by the network dynamics. The parameters regulating these aspects of the physical microenvironment, namely the grid dimensions, the presence of neighbouring cells, and local chemical concentrations, can be customised by the user based on specific assumptions suitable for the system to be modelled (the details for each parameter are discussed in the following sections). These assumptions give rise to certain aspects of physical dimension and spatial competition among cells. Furthermore, cells are also *meta-agents*, namely each cell is itself populated by a community of computational agents, the genes and molecules, which act and interact as a network inside the cell agent.

### Cell mutation profiles

The microC modelling framework enables to input mutation profiles, representing the specific mutations present in each simulated clone. Thus, it might be used to represent the clonal make-up of specific samples, such as 3D mixed-population cellular spheroids. Users may define mutation profiles via an input file uploaded to microC. Such files define the (sub)population of cells where specific gene mutations are present. By default, if not mutation profile is provided, the status of each gene in each cell is initialized randomly as active/inactive. If a mutation profile has been defined, this random initialization is overwritten by the specific mutations. The mutations are introduced in the model as constraints on the gene status, such as a constantly active/inactive status (e.g. constitutively activating or inactivating mutations), or they can be introduced as a change in the rules regulating the gene behaviour (e.g. amplified or conditional behaviour). There is no limitation with respect to the number of mutation profiles that can be simulated simultaneously, or the number of mutated genes. In the extreme case, each cell can have a distinct mutation profile, and multiple mutations can occur in each gene of the encapsulated network.

### Subcellular gene networks

Gene networks are encapsulated within cell agents, and drive their decisions. Although any type of network model may be used in this framework, our current implementation exploits logical Boolean networks. The latter have been shown to preserve key dynamical characteristics of the gene network (26) and they can be designed quickly, without the need for accurate estimates for a large number of parameters. Briefly, the (gene) nodes of such logical networks can have only two states (active or inactive). Nodes may be connected with other nodes via links. All nodes are assigned logical rules that determine the current and future state of the node. We use asynchronous network update, first described by Thomas (27), and widely adopted since in Boolean gene networks. We distinguish between four types of nodes: genes, receptors (input), output nodes, and fate-decision nodes. Genes have both incoming and outgoing links, receptors have only outgoing links, and output and fate-decision nodes have only incoming links. Cell-fate decision nodes have a crucial role in the simulations as they trigger the actions which determine cell behaviour.

In the specific model developed here, we considered as a starting point a previously developed large mitogen-activated protein kinase (MAPK) network (28). This is a Boolean network that has been assembled in a knowledge driven, mechanistic, manner by considering gene-gene relations (e.g. activation/repression) as reported in the published literature. As such, this network it is not specific to a given context, cell line or genotype; however, it is likely to be somewhat biased towards signalling and interactions observed in cancer cell lines, as the pathways modelled are key to the hallmarks of cancer. We then extend this network to introduce a hypoxia responsive module; this new module and the full network can be explored in Figure 3 of the Supplementary Protocol File, and the online documentation.

### Cell status and cell-fate decision rules

An activated cell-fate decision node is associated with a specific action. Our current implementation of the framework includes the following possible actions:

- Proliferation. A copy of the cell is introduced into a neighbouring slot; if all neighbouring slots are occupied, the cell enters the growth arrest state. (This may be modified in future versions of microC in order to allow for different degrees of cellular constraints: from loosely bound cell masses to tightly bound spheroids). The cell-fate decision node is reset and the cell may enter another round of network simulation, and proliferate again, or chose another cell-fate decision. Indefinite proliferation is unlikely to happen due to spatial constrains (all neighbouring slots will be occupied at some point) that will in turn force the cell into a growth arrest state.
- Apoptosis. The cell dies and is removed from the simulation.
- Necrosis. The cells dies; it remains in the simulation and occupies space, but it does not otherwise actively participate.
- Growth Arrest. The subcellular network simulation for a cell in growth arrest is “paused” for a given period of time (3 rounds of simulations in our initial model). Furthermore, the cell interacts with the microenvironment at a reduced rate, for example the consumption rate drops to half of the normal rate.
- No decision. The cell takes no action; the simulation continues until a decision is made.

Actions associated with apoptosis and necrosis are executed immediately after the cell-fate decision has been made, whereas actions associated with proliferation and growth arrest decision are executed shortly after the end of a time frame defined by the user (decision window). During this time frame the network in each cell is simulated and the activation status of cell-fate decision nodes may change.

### A spatially-aware environment

One of the most novel aspects of microC is that cell-microenvironment and cell-cell interactions are modelled, via exchange of diffusible substances between cells, or between a cell and its microenvironment. Importantly, the environment in microC is modelled as computational agent patches. Concentrations of the diffusible substances are transferred through step functions and may activate receptor nodes of the network. Those receptors may in turn trigger an autocrine or paracrine interaction, influencing cells towards specific cell-fate decisions, or the production of certain substances defined by output nodes.

Of note, in microC each 3D voxel is defined as an agent. These voxel or “patch” agents are assigned rules defining their geometry and the interaction with the cells they contain, and how they communicate with other voxel agents. This means that different parts of the environment could be defined by different patches, hence different rules. However, our in initial model and implementation the rules are set as equal for all voxels (e.g. shape, size, number of cell per patch, diffusion). Cell-microenvironment (cell agent-voxel agent) interactions are associated with a list of environmental resources (for example oxygen and growth factors), whereas cell-cell interactions may be the result of user-defined substances, such as cytokines, chemokines, growth factors, and hormones. Diffusible substances are also simulated in microC as agents, and their behaviour is simulated following the diffusion – reaction equation:

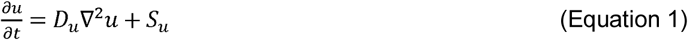

where, *u* is the concentration of the diffusible substance, *D*_*u*_ is the diffusion coefficient of substance *u*, and *S*_*u*_ are sources or sinks of the diffusible substance. The equation is solved numerically using an explicit FTCS (Forward Time Central Space) scheme, with Dirichlet boundary conditions, on a 2D or 3D rectangular lattice. The grid cell size may be adjusted to include 1 (1×1×1) or 27 (3×3×3) cells using the “grid sparsity” parameter.

Cells are modelled as sinks that consume oxygen at a rate proportional to the local oxygen concentration. In particular, oxygen consumption is modelled through the equation:

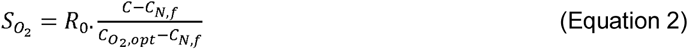

where *R*_*0*_ is the initial consumption rate, *C* is the concentration of oxygen in the specific grid cell, *C*_*N,f*_ is a threshold value that determines the lowest possible oxygen concentration (currently fixed at 80% of the oxygen activation threshold), and finally *C*_*O2,opt*_ is an optimal oxygen concentration, currently set to 0.28mM. The latter two parameters have predetermined values in microC, whereas the initial consumption rate (R_0_) and the oxygen activation threshold may be set by the user. The latter is a precondition that triggers the necrotic cell-fate decisions. We set the necrosis threshold at 0.02mM (23). Cells in growth arrest, consume oxygen at half the initial rate, whereas necrotic cells do not consume oxygen. Parameters such as the diffusion coefficient and the initial/boundary conditions of oxygen concentration can be adjusted as required by the specific application.

Cell-cell interaction is modelled using the diffusion – reaction equation (equation 1), with S_u_ representing the sources and sinks (cells) of hormones, cytokines and any user-defined diffusible substance.

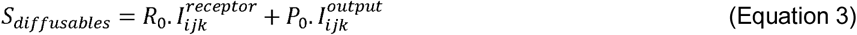

Both the consumption of diffusible substances (consumption rate: R_0_) and the production of these substances by cells (production rate: P_0_) are defined by the user and considered to be constant. Production/consumption of diffusible substances is conditional to the activation status of the corresponding receptor/output node, shown here as a Boolean function I_ijk_ = 0/1, with i,j,k denoting the position of the cell in the 3D lattice.

### Spheroids growth measures

In our growth curves simulations, we use the number of cells as an indicator of the size of the spheroids at any given time, and we have used sphericity to assess their degree of roundness:

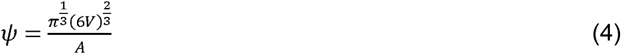

where *Ψ* is sphericity, *V* is the volume of the object and *A* its surface area. The radius of a spheroid is determined at each stage of growth as the average distance between the coordinates of the initial centre point of the simulation, and the outermost cells of the growing spheroids.

### Cloud Implementation

The models presented in this study are accessible via a web interface (http://www.microc.org). This interface also enables modification of the models and input parameters to conduct experiments other than those discussed in this paper. We have prepared a detailed protocol (Supplementary Protocol) that explains how to submit experiments and how to interpret the results. Briefly, the interface allows a user to upload input parameters to set the model (e.g. mutation profiles for the cell populations, inner-cell gene networks, specific values for diffusion, and other kinetic and simulation parameters), it then enables to monitor experimental results in time, and to perform statistical inference on the results. Experiments are specified via a web interface (see Supplementary Protocol), where the user may set a number of numerical parameters via sliders. Specifications of the gene network, mutations, and other parameters can also be uploaded from the same page. The gene network can be specified as a GXL, GraphML or GINML file. GXL is a widely-used XML-based standard exchange format for sharing data; it is a flexible data model that can be used for object-relational data and a wide variety of graphs (29). GraphML is another XML-based, widely-used, data sharing format for graphs (30). GINML is an extension of GXL and can be produced for example by the logical model editor GINsim (14). The web server converts any of the above formats to the GraphML format, and then submits the experiment as a set of jobs to the Advanced Research Computing Cloud (University of Oxford). Experiments are then executed exploiting the NetLogo framework (31). Each node runs 16 repetitions of the experiment on each of its CPU cores. When all the simulations runs have finished, a HTML file containing both the data and JavaScript interactive data visualisations is assembled and a link to the page is sent by email to the user. The URL to the results is automatically generated and is private to the user who can choose to share it. Of note, by implementing microC as a cloud service with a web interface for both submitting experiments and analysing the results we have automated a great deal of technical and tedious work. Files are automatically converted to the necessary formats, pre-defined scripts are used to run the experiments on the cloud, and JavaScript on the results page provides interactive visualisations of the data, data analysis and animations of the cellular model and the gene networks.

## Results

### *In-silico* growth of a heterogeneous population of cells communicating within a three-dimensional microenvironment

We present a modelling framework, microC, which enables simulations of individual cells or group of cells, where each cell contains an inner gene network that simulates the cascade of signalling events occurring in each cell upon stimulation, and determines the cell behaviour (Figure 1). To illustrate microC, we report the different ways in which it can be applied (Figure 1 A-E). Choosing cancer spheroids as a model system, we illustrate how a specific model can be constructed from its individual elements; we carry out simulations under different perturbations (e.g. mutations, nutrients availability); we discuss how our predictions agree or disagree with results from wet-lab studies; and we conduct sensitivity analyses to assess the impact of the different parameter choices on the model predictions.

As initial step, we assessed our ability to import the required gene networks, and confirmed the faithfulness of our ABM encoding of these networks. We then scaled up the simulation by increasing the genetic and microenvironmental heterogeneity in a stepwise manner. Specifically, we considered a population of homogenous pre-cancerous cells growing in a 3D environment; then, we gradually introduced mutations in onco- and tumour suppressor genes (Figure 1A) to study how they affect the multi-cellular growth (Figure 1B). Subsequently, we allowed the resulting clones (carrying different single or multiple mutations) to grow in competition (Figure 1C), and studied the parameters affecting their evolution. Finally, we investigated the additional effects of introducing extrinsic perturbations such as lack of oxygen and presence of growth factors, and enabling cell-microenvironment (Figure 1D) and cell-cell interactions (Figure 1E). We report the parameter values chosen for each experiment in Table S1, while a more detailed description of the parameters may be found in the supplementary protocol and in the online documentation file in microc.org.

#### 1. Evaluation of the dynamics properties of the inner-cell gene networks

Whilst there is a large number of methods, from traditional statistics to emerging deep-learning approaches, which permit accurate prediction of the behaviour of a population of cells given some initial inputs; such methods often constitute a black-box when it comes to interpreting mechanistically the results. They emphasize the predictive ability of the model, with respect to the study of the possible causative mechanisms of the predicted behaviour. In this first example, we show how for each single cell microC enables monitoring of the paths which have followed to arrive at a given prediction. In particular, we show how microC enables studying the dynamical properties of the gene networks within each cell.

Firstly, we verified that microC could correctly import and execute a gene network developed in the GINSim interface (14). This involves to reformat and recodify the network so that it can be executed using our agent-based framework (Figure 2). As we have automated this reformatting and recoding in microC, we needed to check its faithfulness. We asked how intrinsic perturbations, namely mutations commonly observed in cancer cells, and specifically in breast cancer, affect the functioning of the network. To this end, we compared the gene activation profiles across populations of cells carrying different mutations and we determined the differences in the resulting fate of the cell, focusing on the three possible decisions of proliferation, growth arrest and apoptosis, and then necrosis when we introduce oxygen diffusion. This first simulations showed that we could reproduce the stable configurations (or stable states) which were previously reported for this network in the published stable-state analysis (Figure 2A and Figure S1A). This illustrates the ability of microC to execute faithfully a previously defined logical network using a ABM approach. Since our protocols are standard, this opens the technical possibility of reusing any such networks developed using the same standard formats.

**Figure 2.**
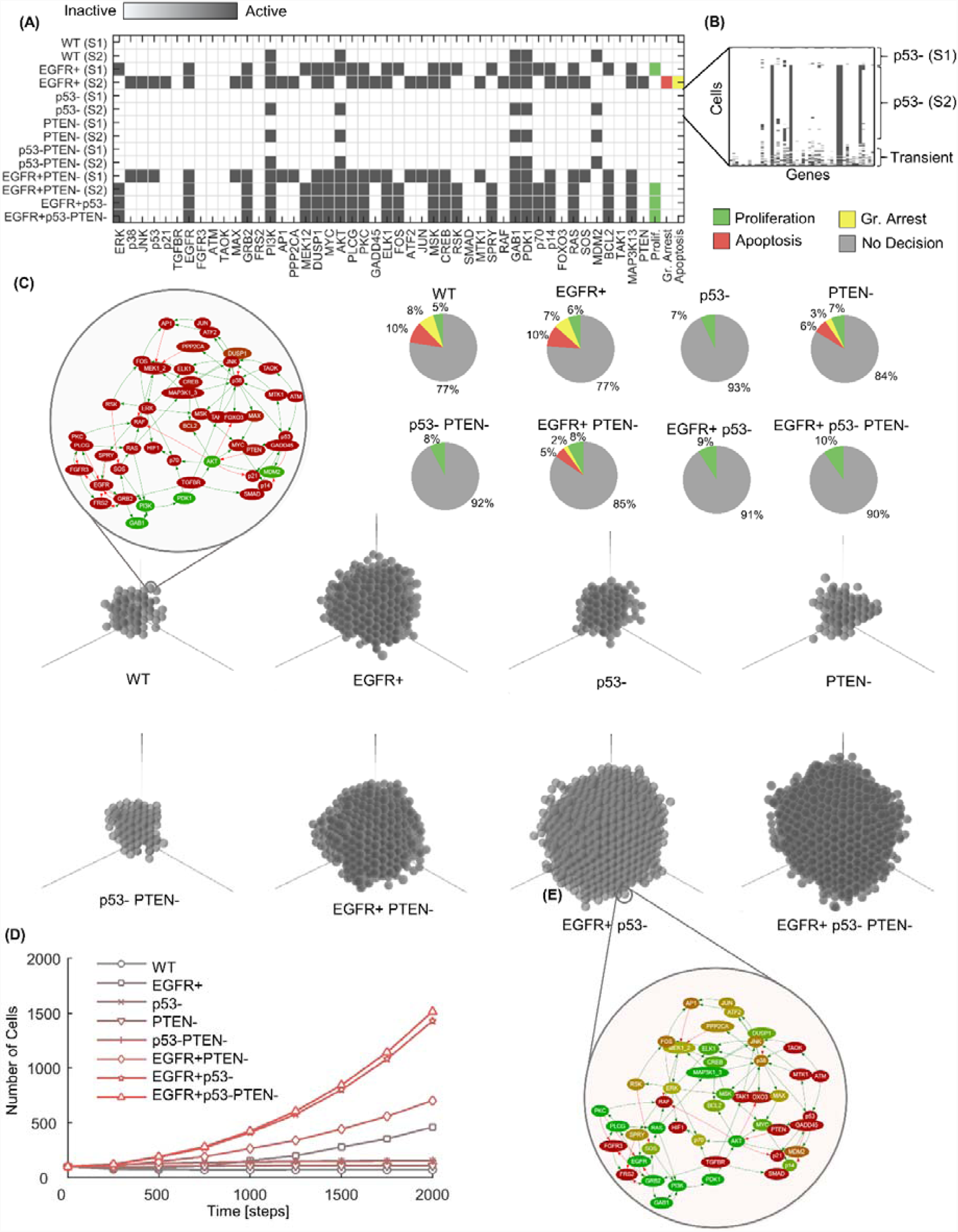
microC simulation of 3D tumour multi-cellular spheroids reveals growth advantage provided by co-occurring mutations. A. Activation status of genes in the MAPK network for the stable states. The states are labelled with the appropriate colour (Green: Proliferation, Red: Apoptosis, Yellow: Growth Arrest) depending on the cell-fate node activated. Cells with no specific decision, are not labelled with any colour. B. Detailed activation status for the clone with p53 loss-of-function mutation (p53-). Columns represent genes and rows cells (initial population: 100 cells, repeats: 100, length of simulation: 5000 steps). Pie charts represent probability profiles for cell-fate decisions for each clone. C. Typical examples of spheroids grown in a 3D environment, and MAPK network for two of the clones (upper left for WT cells, which grow very little, and right below for EGFR+p53-cells, one of the most aggressive clones). Circle represent the cell boundary, inside network nodes represent gene products (colour represents activation status, red is inactive and green active); edges carry the information on the gene-gene interaction (here green is activation and red is inhibition). D. Clonal growth curves (initial population: 100 cells, repeats: 100, length of simulation: 2000 steps).

In addition to the published results, and thanks to the ABM approach, we could also evaluate the probability of occurrence and temporal profile for cell-fate decisions by performing hundreds of replicated simulations (Figures 2B, Figure S1B, S2). This reveals that not all stable-states are equally likely to occur, and it uncovers transient states that would remain undetected in a standard stable state analysis. For example, we found that there was a contained but measurable (7%) probability of proliferation for cells with a single loss-of-function mutation in p53 (p53-) (Figure 2B). Often transient states have been dismissed as non-biologically relevant. However, if and when they are, or not, is neither a trivial nor fully answered question. Thus, to detect such states in a network analysis and highlight their potential impact on the different fate decisions, is important in order to start to address their relevance.

#### 2. Effect of mutations on gene network dynamics and multi-cellular growth

The range of abilities, or hallmarks, that a cell needs to acquire in order to progress to become a cancer cell have been extensively described, and the role of somatic mutations in such processes is well documented (9). However, the chain of events from a single mutation occurring in a normal cell to the acquisition of such hallmarks, and then to the occurrence and progression of cancer, is extremely complex and not fully understood. Here, we used microC to dissect such question by asking how the occurrence of single and multiple mutations might impact on the behaviour of signalling networks, and how this results in different patterns of single-cell and multi-cellular growth.

We focused on mutations frequently occurring across cancer types, namely loss-of-function/inactivating mutations in the well-known tumour suppressor genes p53 and PTEN, and the activation of the known cancer driver EGFR. We then monitor the 3D growth patterns of cells where these mutations were introduced as single mutations (EGFR activating mutation, EGFR+; p53 loss-of-function, P53-; or PTEN inactivation, PTEN-), or in combination of two or three. We compare the resulting growth curves with those predicted using a wild type (WT) simulation, namely cells not carrying any of these mutations. Of note, for these initial simulations we consider a media with no growth factors nor other added nutrients, simulating a condition of cell starvation (see Table S1 for all parameter choices).

In these conditions, our model predicts that clones carrying multiple mutations are characterised by a significantly more aggressive phenotype with faster growth with respect to clones carrying single mutations in these genes (Figure 2C and 2D, and Figure S3). Furthermore, our model predicts that activated EGFR signalling is a determinant for rapid growth under starved conditions. Specifically, in our simulations all EGFR+ clones (EGFR+, EGFR+PTEN-, EGFR+p53-, EGFR+p53-PTEN-) all exhibit initial exponential growth while clones with no activated EGFR signalling did not grow or grew at a much slower rate (Figure 2D).

To assess the usefulness and accuracy of our predictions we considered experimental data obtained using cells where mutations in EGFR, p53 and PTEN have been induced either alone or in combination. We asked how an initial general model, not specifically built for any cancer types, but built in a mechanistic fashion using hundreds of independent publications on gene interactions could provide useful predictions with respect to published studies not used in the model building. Specifically, we considered results from a study using MCF10A, an immortal though not tumorigenic mammary tumour cell line. This model has been shown to express markers which are associated with the basal-epithelial phenotype, but it does not carry the mutations frequently observed in this breast cancer type. In these cells, the effect of PTEN deletion, p53 loss-of-function mutation and EGFR activating mutations on growth and colony formation has been previously measured under conditions of starvation (32, 33). As shown by Pires et al (33), these mutations as individual oncogenes could not stimulate growth in 3D culture in soft agar, whilst the double mutants showed increased growth, and the triple mutant grew significantly more rapidly and formed significantly more colonies than either of the matched double mutants. These experimental results agrees with our simulations. We though observed also a discrepancy between Pires et al experiments and our predictions as in their hands PTEN as individual oncogene could stimulate 2D growth, but not 3D growth, of MCF10A cells in the absence of exogenous growth factors. In our predictions we did not observe any significant difference between the 2D and 3D growth of the PTEN clones.

We then compare the sphericity as further geometrical property of the spheroids growth in a 3D environment, other than simply size in time. Sphericity is a measure of how near an object is to a perfect sphere (see Methods). This showed that spheroids formed by the most aggressive clones (higher proliferations rates), namely EGFR+p53- and EGFR+p53-PTEN-, not only grew faster but they also grew in a more symmetrical manner, with higher sphericity than other clones (Figure S4). This is understandable from a geometrical point of view, as in our model the clones which tend to have proliferation as their main action, rather than apoptosis or growth arrest, can make best use of the space around them. However, the accuracy of this prediction would need to be confirmed. In this respect, the difficulty of growing some of the less aggressive cell lines means that there is not published evidence on these morphological aspects, and they are hard to assess in an experimental context. However, this is an intriguing result which suggest that morphological characteristics might be a useful aspect to consider in future studies linking the way a spheroid grows with its clonal composition.

Finally, we compared the relative doubling time of the different clones. We observed that the doubling time for the EGFR+ clones (EGFR+, EGFR+PTEN-, EGFR+p53-, EGFR+p53-PTEN-) ranged between 4.1 – 6.9 rounds of simulation, thus it was relatively short, whilst the rest of the clones did not have the potential to double. These results agree well with results reported by an independent study (not used to train the model) for pre-malignant MCF10A breast epithelial cells carrying EGFR activating mutations, with respect to wild type MCF10A cells (34). Specifically, EGFR mutant cells showed a relatively short doubling time (20-22 hours) irrespective of the presence of EGF in the media, whilst cells not carrying a EFGR activating mutation did not reach their doubling time, unless stimulated by EGF.

#### 3. Scrutiny of clonal evolution paths: significance of the order at which mutations occurs

In the previous examples, we considered the co-existence of multiple mutations. This reflects reasonably well experimental conditions where mutations are artificially introduced, but it might not represent clonal evolution in real tumours. Scrutiny of clonal evolution paths is an emerging field of research, however this has typically been done without taking into account the signalling and context in which mutations occur. In this example, we ask how the order in which mutations occur affects both individual cells and the overall multi-cellular growth.

Focusing on the aggressive EGFR+p53-PTEN-clone (Figure 3A), we examine the six possible evolution paths for this clone (Figure 3A-B). This revealed that the order in which mutations occur has a significant effect on a spheroid’s growth (Figure 3C). Specifically, the time of occurrence of the EGFR+ mutation was crucial: clones that acquire this mutation before the loss of a tumour suppressor resulted in larger spheroids, followed by those clones that acquire the EGFR+ mutation, and finally those that acquire the EGFR+ mutation (Figure 3C).

**Figure 3.**
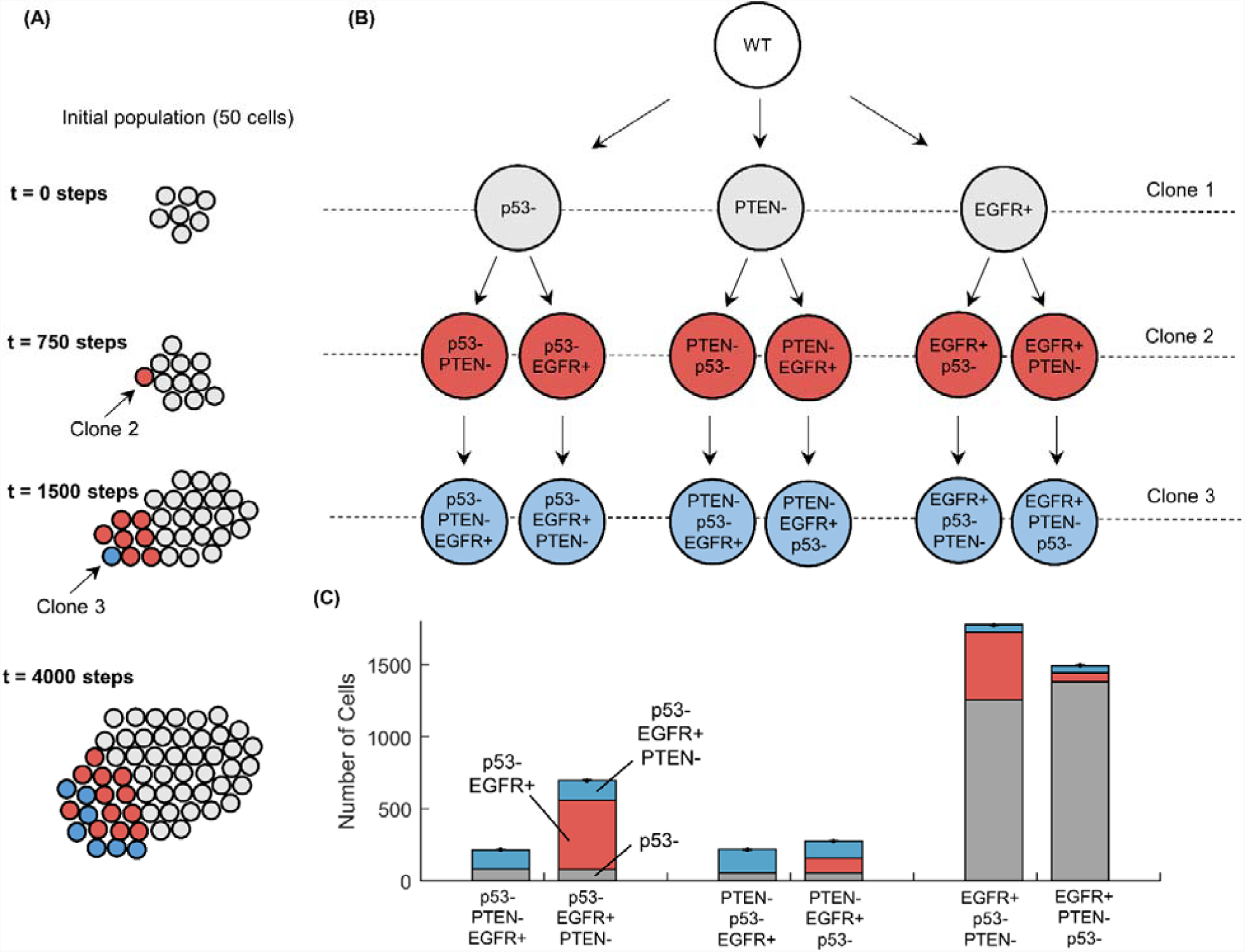
Paths of clonal evolution: the order in which mutations occur affects spheroids’ growth. A. Experimental setup. The experiments start with 50 cells of the same clone (Clone 1, shown as grey spheres). At time point t = 750, we introduce a mutation to one cell (Clone 2, shown as red spheres), and at time point t = 1500 we introduce an additional mutation to a cell of type Clone 2 (that we call Clone 3, shown as blue spheres). We extend the experiment up to 4000 simulation steps. B. There are six potential combinations in which the three mutations (p53-, PTEN-, EGFR+) can occur. C. The total and clonal subpopulations are shown in the stacked bar chart. Bars are averages of 100 repeats and error bars represent the standard error.

These results confirm first that cancer can occur from multiple evolutionary paths, but they also suggest that a proliferation stimulus, namely EGFR activation, followed by the loss of a tumour suppressor, p53 loss-of-function, and then PTEN loss-of-function, results in the most rapid evolution. This appears to support the preponderance of experimental data indicating that p53 mutations is a relatively late, rather than cancer initiating, event in a number of cancers (see e.g. (35)). However, the evidence on this point is contrasting, and it is well known that p53 is affected by multiple mutations, with different functional implications (for a review see e.g. (36)), so further studies are needed to include such differences. Importantly, and differently from other approaches to study clonal evolution, microC enables exploration of how differences in growth rates between cells carrying different mutations might result from the underlying characteristics of the network (Figure 2 and S5). For example, EGFR activation directly affects the status of a large group of genes including *ELK1, CREB, MYC* and *RAS* that promote proliferation and block apoptosis (Figure S6), so to acquire this mutation early would be very advantageous.

Finally, growth rates depend also on dynamical network parameters, such as the speed of the specific intercellular processes. We performed a thorough sensitivity analysis by changing the value of the parameters, starting from values suggested in the literature and expanding the range far from this initial choice (see Methods and Supplementary Material, and Figure S7-8). This identified the temporal decision window as one of the critical parameters, which is tightly linked with the temporal ratio between intracellular to intercellular processes. This analysis shows that changes in this parameter can affect the resulting growth rate, and the effect is different when different mutations are considered. As expected, variation in the decision window were reflected by a different number of cells associated with a given cell fate, as more cells could reach a decision when longer windows where allowed. Small decision windows (below 100 steps) showed smaller differences between the growth curves of the different clones; however at this time only a small number of cells will have reached a decision (Figure S2), thus the predictions are based on a very small number of cells. For large decision windows values, the predictions started to converge and the choice on the decision window affected the absolute number of cells, but not the relative populations and the ranking of the clones with respect to their ability to proliferate was maintained (Figure S7). This was also reflected in the activation/inactivation profiles for the different states in the different clones, where a value of 100 was the optimal choice in order to preserve dynamical behaviour of the model, while minimizing computational intensity of the simulations (Figure S8).

#### 4. Emerging competition patterns impact the growth of multi-clonal cell populations

Next, we asked how competition between different clones, grown together in a 3D multi-cellular spheroid, affects the 3D growth dynamics. To this end, we introduce multiple clones in the same environment (Figure 4A and 4B), and compare the growth curves of the resulting spheroids (Figure 4C), and their final size (Figure 4D), with those observed when the same clones were grown in isolation, that is in single-clone spheroids.

**Figure 4.**
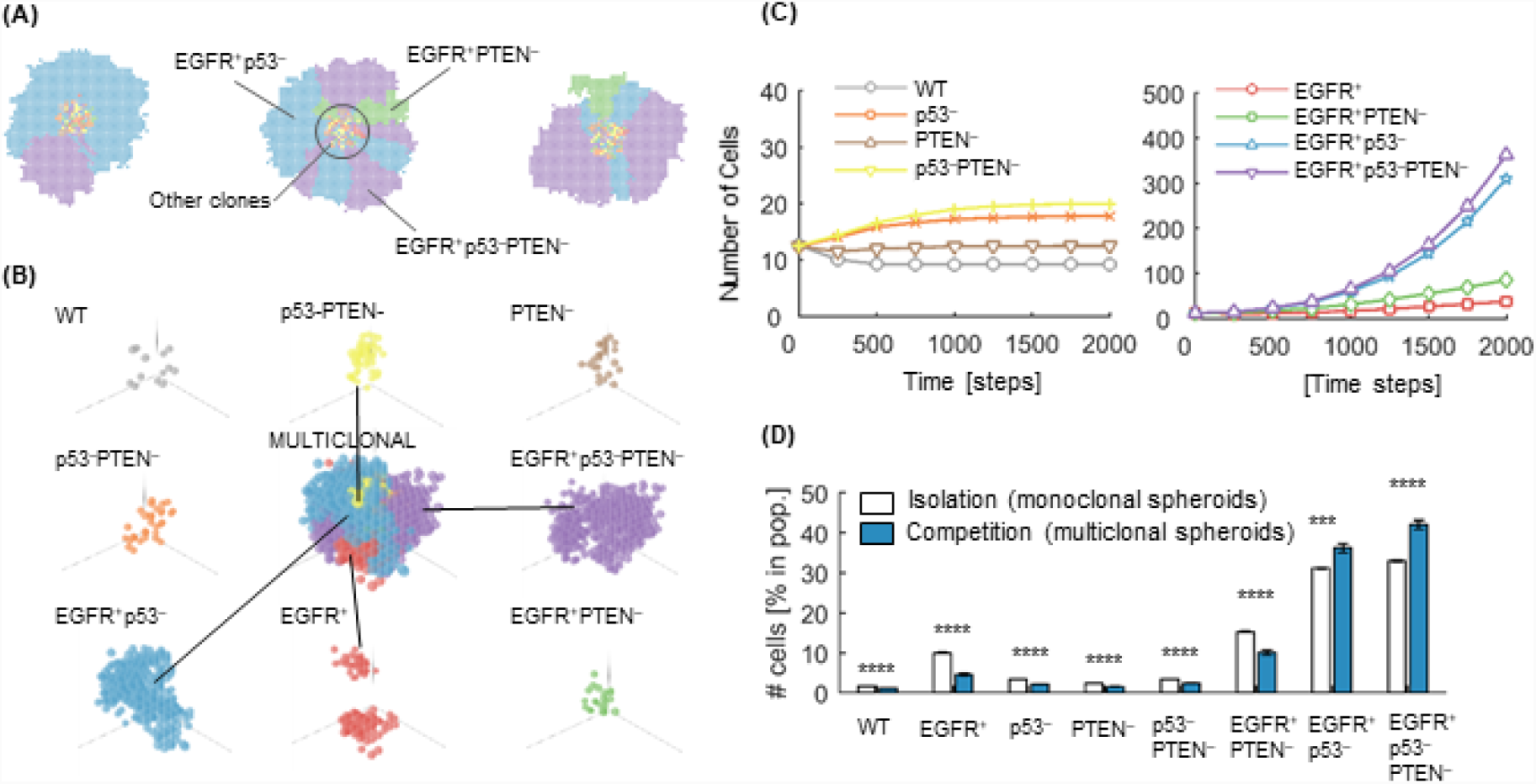
Growth of multi-clonal spheroids is affected by, and in turns affects, the competition between the clones. A-B. Cells with different mutation profiles (colours corresponds throughout) are grown within the same virtual microenvironment. At the end of the simulation each cell type shows a distinct growth pattern. Here patterns are shown in 2D and 3D. Figure B shows a spatial decomposition of a multiclonal spheroid to demonstrate the distribution of the different clonal populations within the spheroid. C. Growth curves of clones under competition in 3D simulations. D. Relative clonal population (over total population of cells) under isolation and competition in 3D simulations. Results are averages of 100 repeats of an initial population of 100 cells. Under competition the initial population was shared in equal parts among the eight clones. Error bars in bar charts represent standard error. * indicates Kruskal–Wallis p<0.05, ** p<0.01, *** p<0.001, **** p<0.0001.

In the multi-clonal simulations, we reveal that clones with aggressive phenotypes systematically take over the free surface area of the spheroids, thus restricting the rest of the clones in the central parts of the spheroid. This is particularly evident in the 2D geometry (Figure 4A). This is a striking finding and it implies that the population of the aggressive phenotypes not only increases rapidly against the rest of the clones due to faster growth, but it also creates a physical barrier to for any further proliferation of the inner, slower growing, population.

This effect significantly affects both the growth curves (Figure 4C) and the final population fractions (Figure 4D) achieved by the same clones grown in competition and in isolation. The most aggressive clones (EGFR+p53- and EGFR+p53-PTEN-) increase their presence by 16.7% (EGFR+p53-, from 31.1% of total population in isolation to a 36.3% in competition), and by 27.3% (EGFR+p53-PTEN-, from 33.0% of total population in isolation to a 42.0% in competition). The rest of the clonal populations decrease by 28-54%. The population of cells that decrease the most (by 54.5%) are the EGFR+ clones.

Interestingly, these findings can be examined in light of the proliferation rates and the dynamical properties of the different clones (Figure 2B). In particular, EGFR+p53- and EGFR+p53-PTEN-clones, are the only ones characterized by a single possible outcome for the cell fate decision, which is proliferation. This is a strong advantage when competing with clones that are characterized by multiple decision outcomes. For example, EGFR+ clones have a higher probability of undergoing apoptosis or growth arrest, which constitutes a strong disadvantage under direct competition since apoptotic events free space that are likely to be rapidly occupied by more aggressive clones.

#### 5. Interaction between the genotypes and the microenvironment affects multi-cellular growth and clonal selection

Although the mechanisms are not yet fully elucidated, increasing evidence points to a key role of the microenvironment in clonal selection, hence multi-cellular growth of heterogeneous populations. To illustrate how microC can address this, we focused on hypoxia simulations as this is one of the major microenvironmental differences between cancer and normal tissue (37-40). Specifically, we aim to study the formation of necrotic cores in larger spheroids (Figure 5A-B), and their growth under artificially uniform well-oxygenated conditions (our *Control* configuration) with respect to growth when oxygen consumption and diffusion are enabled (we refer to this as the *Hypoxia* configuration) (see also Methods section). To achieve this, we consider environmental agents that simulate the diffusion of oxygen in the microenvironment and have the ability to trigger the hypoxia responsive module (see Methods); we also introduce a new cell fate decision, necrosis, as response to extremely low oxygen concentrations (0.02mM O_2_,). We then simulate this model in our control and hypoxia configurations.

**Figure 5.**
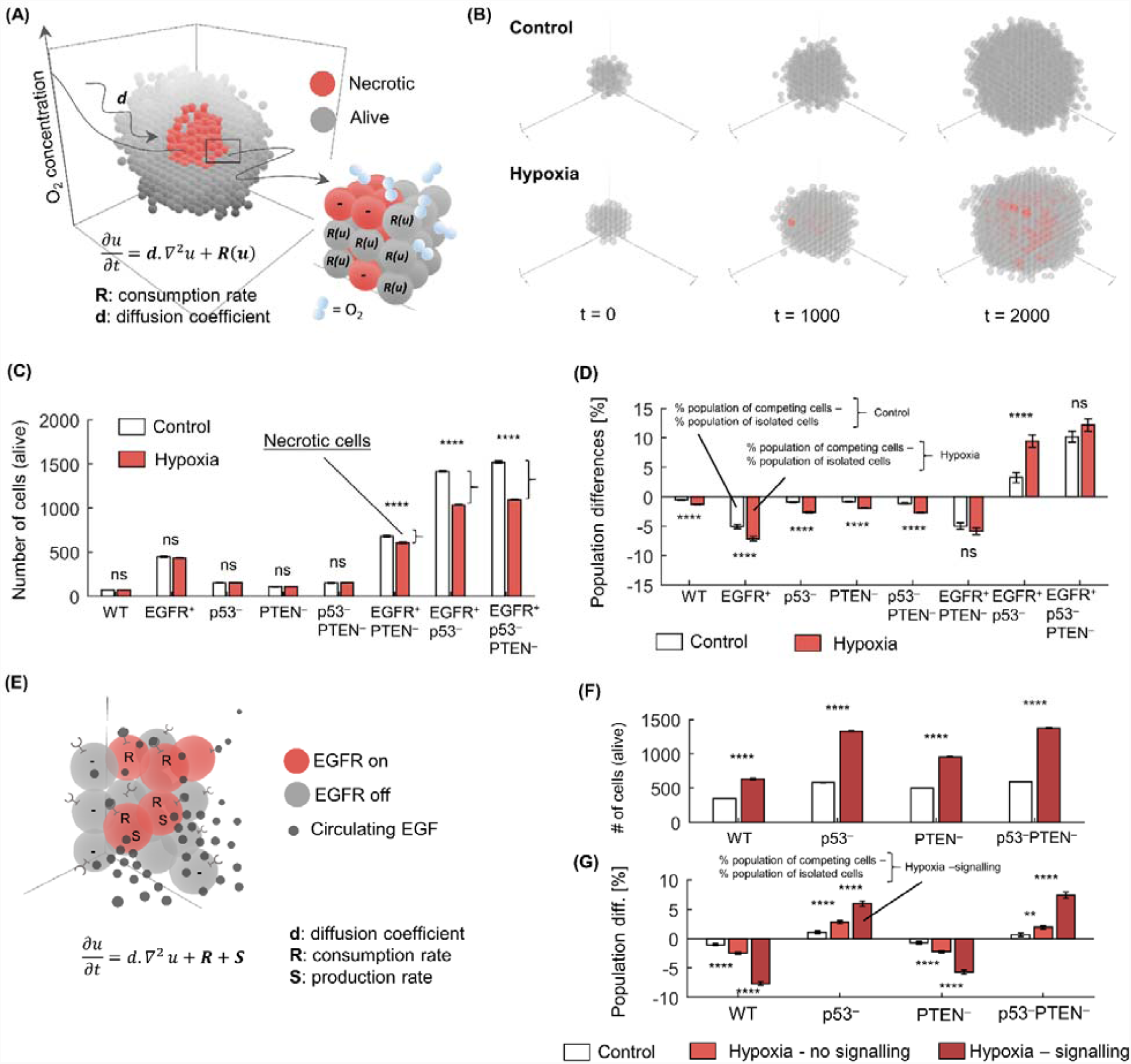
The effect of a hypoxic microenvironment and cell-cell signalling on multi-cellular growth. A. Necrotic core within a spheroid as a result of oxygen consumption by cells. Necrotic cells do not consume oxygen (-), in contrast to living cells (R: Oxygen consumption rate). Oxygen diffusion is simulated by the diffusion-reaction equation (Equation 1, in Materials and Methods). B. Spheroids growing under artificially well-oxygenated conditions (*Control*, initial and boundary condition: 0.28 mM O_2_) and an environment with oxygen level drop in the inner layer of the 3D spheroid due to diffusion (*Hypoxia*, initial and boundary condition: 0.04 mM O_2_). C. Growth under the well-oxygenated (Control) condition and the hypoxic condition (Hypoxia), for eight clones. EGFR+ clones develop necrotic cores under the hypoxic condition (differences between white and red bars (right braces) show the number of necrotic cells). D. Differences in clonal population fractions (in-silico living cells) between cells grown isolated and in competition to each other, under well-oxygenated condition (Control) and the hypoxic condition (Hypoxia). Bars are averages of 100 repeats (initial population: 100 cells, experiment length: 2000 temporal steps, R_O2_ = 5.0e-3 mM. s^−1^, D_O2_ = 1.0e-9 m. s^−1^), and error bars represent standard error. * indicates Kruskal–Wallis p<0.05, ** p<0.01, *** p<0.001, **** p<0.0001. E. EGF signalling between cells. Red-coloured cells have activated EGF receptors, whereas grey-coloured cells have inactivated EGF receptors (junctions on cells). EGF molecules (small, black spheres) are produced (S: EGF production rate) by cells and may be consumed (R: EGF consumption rate) by activating the EGF receptor of the same or a neighbouring cell (spheres attached to the EGF receptors), thus promoting proliferation. Cell – cell interaction is simulated by the diffusion – reaction equation (see Material and Methods). F. Spheroids grown with disabled EGF signalling (*Hypoxia - no signalling*), and with enabled EGF signalling (*Hypoxia – signalling*, 0.04mM O_2_, ACT_EGF_ = 5.0e-4 (^+^).m^3^, R_EGF_ = 5.0e-4 (^+^).m^3^.s^−1^, R_O2_ and d_O2_ as above), initial population: 500 cells, experiment length: 2000 temporal steps. The initial and boundary configuration under Hypoxia is 0.04 mM O_2_. Bars are averages of 100 repeats. Error bars represent standard error. G. Relative clonal populations in isolation and competition, under different oxygen configurations (Control (Normoxia), Hypoxia – no signalling, and Hypoxia – signalling). Percentages are normalised for each condition to the number of living cells at the end of the simulation across the four cell types. Bar charts are averages of 100 repeats (initial population 500 cells, 2000 steps). Error bars represent standard error. * indicates Kruskal–Wallis p<0.05, ** p<0.01, *** p<0.001, **** p<0.0001. (^+^) fraction of the EGF production rate.

Of note, in our network (Supplementary Protocol Figure 3) the activation of the hypoxia responsive module causes production of EGF by the cell, which can then be sensed by the neighbouring cells and activate EGFR. Thus, the normoxia simulation is carried out in the absence of EGF (no production and no prior presence in the environment), whilst EGF is present in the hypoxia configuration (produced by the cells). Of note, we can modify this element and either dissociate EGF from the receptor (so EGF present but not sensed, representing for example a drug treatment inactivating the receptor) or we can decide not to simulate EGF (so EGF not in the environment, representing a knock-out of the EGF gene). This allows us to check the effect of introducing signalling between cells in the model, and conversely the effect of interfering with this signalling (Figure S3).

The first simulations described below are carried out with this signalling switched off and simply assess the diffusion parameters and occurrence of necrosis. In the next session we will evaluate the EGFR and cell-cell signalling component. Interestingly, we observe that EGFR+ clones are the only ones with the potential to outgrow a critical spheroid size that triggers necrosis. For a configuration setup with initial consumption rate 0.005mM.s^−1^ and diffusion coefficient 10^−9^ m^2^.s, the necrotic core that was formed ranged between 20 – 360 cells (Figure 5C) in agreement with previous reports (41). Accordingly, spheroids with necrotic cores had on average 4.8% - 23.6% less living cells than spheroids without necrotic cores.

We then study the combined effect of hypoxia (as defined above), and clonal competition (see section 3). Namely, we introduce all eight clones in the simulation environment and compared growth in isolation and competition, under either hypoxic and control conditions (Figure S9). Overall, our simulations showed that spheroids where hypoxia was enabled had reduced clonal diversity (Figure 5D). In particular, we noticed that the EGFR+p53- and EGFR+p53-PTEN-clones increased their presence in the total population. This was additional to the initial enrichment of this population due to competition (white bar series in Figure 5D). On the contrary, the rest of the clones decreased their presence in the spheroids by an average of 4 - 5%. This selection pressure, and consequent reduction of clonal diversity due to hypoxia can be explained by the spatial distribution of clones (Figure 4A). Less aggressive clones are more likely to be segregated to the central part of the spheroids, that eventually will become necrotic under hypoxia. Of note, in this example we have not introduced possible mutations which might occur as a result of hypoxia, which could impact on the observed clonal diversity.

Finally, we perform a sensitivity analysis on the parameters involved in the diffusion-reaction equation (Equation 1), namely we change the oxygen consumption rate and the diffusion coefficient to one of a broad range of values (Figure S10). We found that the consumption rate was the only parameter which, when changed, significantly affects oxygen concentration (Figure S10A), which may impact spheroid growth indirectly by triggering necrosis. This prediction is in agreement with previous evidence showing oxygen consumption to be the most critical kinetic parameter (41).

#### 6. Impact of cell-cell signalling on multi-cellular growth in a heterogeneous microenvironment

Our final example asks how accounting for cell-cell interaction affects the growth of spheroids. This experiment highlights one of the most innovative aspects of microC. As cell signalling and its relevance for normal development and disease are increasingly understood, a framework which enables such modelling, and study of the consequences of such signalling on cell behaviour, is correspondingly advantageous.

We study EGF signalling (Figure 5E), a response triggered by many stress factors, including hypoxia. We compare growth under two experimental conditions: Oxygen diffusion simulations with disabled EGF signalling (which we call *Hypoxia - no signalling*, same as *Hypoxia* in section 5), and Oxygen diffusion simulations with enabled EGF signalling (which we call *Hypoxia - signalling*), thus where the cells are producing growth factors which are sensed by the nearby cells. For this experiment we used clones that did not constitutively activated EGFR.

We observed a significant increase in the population of cells under the *Hypoxia – signalling* condition. This increase ranged between 130 - 870 living cells, that is 26 – 174% of the spheroid size (Figure 5F). These effects were consistent with the aggressiveness of the clones that we have seen in the previous examples; namely, p53- and p53-PTEN-clonal populations increased significantly more than the PTEN- and WT clones under hypoxia when EGF signalling was accounted for. Interestingly, enabling EGF signalling between cells further increased the tendency of hypoxia to reduce clonal diversity (Figure 5D and 5G). This reveals that the reduction in clonal diversity attributable to central necrosis (Figure 5D), is further increased due to the different proliferation rates of clones which are sensing EGF released under hypoxia.

A sensitivity analysis of the parameters determining the strength and the length of the EGF response, namely the activation threshold of the stimulus receptor and consumption rate of the growth factor, showed that low activation threshold values are more likely to activate the EGF receptor for any given EGF concentration above the threshold value, whereas lower consumption rates of EGF are more likely to activate the EGF receptor for longer (Figure S10B). Both events increase the probability of proliferation, that in turn affects the size of the spheroid (Figure S10C).

## Discussion

To respond to the need to deepen understanding of the intricate relationship between cell genotype and phenotype, we have developed microC, which combines a powerful stochastic method, ABM, and gene networks modelling, into a novel framework for in-silico experimentation. Specifically, microC addresses the challenge of modelling and simulating the dynamics and evolution of a heterogeneous population of cells within a changing microenvironment. ABM is increasingly used to simulate the dynamics and evolution of complex systems in applications ranging from engineering to ecology (42-45); this approach makes microC a naturally multiscale framework, enabling assessment of both multi-cellular systems and individual cells, and their physical and molecular properties.

One of the most innovative aspects of microC is that it substantially extends the capacity of ABM simulations of living cells, considering each individual cell as a *meta-agent*. Each cell is considered as a community of computational agents, the genes and molecules, acting and interacting within each cell. This enables the user to modify and/or replace gene networks in the cells, to define new constrains representing different types of mutations in different cells or in the same cell, and to customize cell-cell and cell-microenvironment interaction parameters. As a consequence, the proposed framework is naturally suited to study how the behaviour of a specific perturbation, such as a mutation, occurring in individual cells within a three dimensional, dynamic microenvironment, affects the collective behaviour of other, similar or different, cells and the whole system. This reveals in some cases unexpected patterns of collective behaviour, not *a-priori* defined in the model, and which could not be predicted by observing the individual elements. Instead, there are emerging properties of the system as a whole and affect the evolution of the cell population and the behaviour of the single elements in return.

microC comes with an example signalling network that can be used as it is, be modified to model in more detail specific pathways, or be completely replaced with a new one by the user so that it is more specific to system studied. Using this signalling network, we illustrated microC features through the study of growth patterns of 3D *in-silico* spheroids, affected by perturbation of intrinsic (mutations) and extrinsic (oxygen and growth factor availability) factors.

In agreement with experimental results obtained by inducing mutations in the MCF10A pre-cancer model, we demonstrated that clones carrying concomitant mutations in the well-known tumour suppressor genes p53 and PTEN, together with the activation of the known cancer driver EGFR, had a growth advantage with respect to clones carrying single mutations, or combinations of any two mutations. We also predict that the order in which these mutations are acquired can have a significant impact on the spheroid growth rate and final size. Both of these results held true both when the clones were grown in isolation and when they were allowed to compete with other clones. We also showed that this effect increased under more extreme microenvironmental conditions, namely lack of oxygen. Interestingly, we observed that morphological characteristics of spheroids of the same clone varied considerably and may be a source for significant variability in *in-vitro* experiments (46). We show how the dynamics of a gene network which is specified by the genotype and microenvironmental characteristics, affects the proliferation rate, which in turn has a significant impact on the overall spheroids’ size and shape, irrespective of other elements such as adhesion forces which are long known to affect morphology (47).

Finally, we observed expected emerging properties, such as the formation of necrotic cores in the larger spheroids, and new unpredicted emerging properties, such as the effect of hypoxia and EGF signalling on clonal diversity of the spheroids. Importantly, these properties were not encoded at the cellular level, but rose from the cell agents competition for nutrients, space and cellular interaction.

Notwithstanding the modelling possibilities that this novel framework opens, it is important to recognise that the current implementation of the methodology has some limitations. Firstly, although microC accounts for certain important aspects of physical modelling such as the three-dimensionality, the presence of neighbouring cells, and local concentrations of chemicals and molecules, it does not account in its current implementation for physical factors such as matrix porosity, stiffness, and topographical cues. Given the importance of modelling these factors (48, 49), future extension of microC will need to include them. On the other hand, we chose to model the cellular environment using ABM; namely, our 3D voxels are themselves agents whose shape and behaviour is defined by rules (see Methods). These rules can be different for different parts of the environment and can change in time, facilitating the dynamic modelling of physical factors using this framework.

A second limitation of our initial model is that it considers only some possible actions for the cells (e.g. proliferation, growth arrest, apoptosis), and it does not consider for example an important action such as invasion. Whilst enhanced overall proliferation can provide an indication of invasion, and it has has been shown to correlate with migration and invasion capability in multiple cancer cell lines (50), whether specific cells in a spheroid divide or migrate out of the spheroid is an important aspect to address (50, 51). Thus, we do foresee this as an important future development, and our choice of considering cells as independent computational agents in a ABM framework is ideal and greatly facilitates addition of cell actions such as migration and invasion.

Finally, the use of Boolean networks to represent gene expression and activation is a simplification that can be reasonably applied in the case of loss-of-function perturbations, where a gene is either expressed or not, active or inactive. However, this might not accurately represent subtle changes in expression with different biological implications, or mutations which change the gene function instead of simply inactivating the gene. Although it is possible to introduce multiple nodes for a single gene, each one representing a different state, it is natural to extend the current Boolean framework into a logical framework with any number of states covering different levels of expression, or different mutations.

In summary, in this paper we presented for the first time, and assessed the capabilities of a novel modelling framework that links genotype with phenotype, via gene networks and signalling pathways. We have provided a number of examples of using this framework, illustrating strengths and limitations of the current implementation. Importantly, this framework not only enables a broad range and new types of modelling studies, but it also delivers a microenvironment for in-silico experimentation built using well-recognised formats and shared standards, thus enabling widely used model representations. This provides an environment for prediction, experimentation and reasoning using existing gene networks and cell models, as well as a powerful starting point for the development of new ones.

## ACKNOWLEDGEMENTS

The authors would like to acknowledge the use of the University of Oxford Advanced Research Computing (ARC) facility in carrying out this work. http://dx.doi.org/10.5281/zenodo.22558. We also like to thank L Grieco for his insights in the MAPK gene network, and Prof A Harris and Prof JM Brady for their comments and suggestions during the manuscript preparation.

## AUTHORS CONTRIBUTION

FMB conceived the idea, designed and supervised the study. DV and FMB developed the models and interpreted results. DV and KK implemented the modelling interface. MH and RW implemented the interactive 3D visualization. DV and FMB co-wrote the manuscript, with input from all authors.

## References

1. Genomes Project C, Auton A, Brooks LD, Durbin RM, Garrison EP, Kang HM, et al. A global reference for human genetic variation. Nature. 2015;526(7571):68–74.

2. Shalem O, Sanjana NE, Zhang F. High-throughput functional genomics using CRISPR-Cas9. Nat Rev Genet. 2015;16(5):299–311.

3. Ritchie MD, Holzinger ER, Li R, Pendergrass SA, Kim D. Methods of integrating data to uncover genotype-phenotype interactions. Nat Rev Genet. 2015;16(2):85–97.

4. Lehner B. Genotype to phenotype: lessons from model organisms for human genetics. Nat Rev Genet. 2013;14(3):168–78.

5. Yi S, Lin S, Li Y, Zhao W, Mills GB, Sahni N. Functional variomics and network perturbation: connecting genotype to phenotype in cancer. Nat Rev Genet. 2017.

6. Gilman SR, Chang J, Xu B, Bawa TS, Gogos JA, Karayiorgou M, et al. Diverse types of genetic variation converge on functional gene networks involved in schizophrenia. Nat Neurosci. 2012;15(12):1723–8.

7. Girard SL, Dion PA, Rouleau GA. Schizophrenia genetics: putting all the pieces together. Curr Neurol Neurosci Rep. 2012;12(3):261–6.

8. Hanahan D, Weinberg RA. The hallmarks of cancer. Cell. 2000;100(1):57–70.

9. Hanahan D, Weinberg RA. Hallmarks of cancer: the next generation. Cell. 2011;144(5):646–74.

10. Karlebach G, Shamir R. Modelling and analysis of gene regulatory networks. Nat Rev Mol Cell Biol. 2008;9(10):770–80.

11. Le Novere N. Quantitative and logic modelling of molecular and gene networks. Nat Rev Genet. 2015;16(3):146–58.

12. Molinelli EJ, Korkut A, Wang W, Miller ML, Gauthier NP, Jing X, et al. Perturbation biology: inferring signaling networks in cellular systems. PLoS Comput Biol. 2013;9(12):e1003290.

13. Haydarlou R, Jacobsen A, Bonzanni N, Feenstra KA, Abeln S, Heringa J. BioASF: a framework for automatically generating executable pathway models specified in BioPAX. Bioinformatics. 2016;32(12):i60–i9.

14. Chaouiya C, Naldi A, Thieffry D. Logical modelling of gene regulatory networks with GINsim. Methods Mol Biol. 2012;804:463–79.

15. Hoops S, Sahle S, Gauges R, Lee C, Pahle J, Simus N, et al. COPASI--a COmplex PAthway SImulator. Bioinformatics. 2006;22(24):3067–74.

16. Shannon P, Markiel A, Ozier O, Baliga NS, Wang JT, Ramage D, et al. Cytoscape: a software environment for integrated models of biomolecular interaction networks. Genome Res. 2003;13(11):2498–504.

17. Hanahan D, Coussens LM. Accessories to the crime: functions of cells recruited to the tumor microenvironment. Cancer Cell. 2012;21(3):309–22.

18. Polyak K, Haviv I, Campbell IG. Co-evolution of tumor cells and their microenvironment. Trends Genet. 2009;25(1):30–8.

19. Orgogozo V, Morizot B, Martin A. The differential view of genotype-phenotype relationships. Front Genet. 2015;6:179.

20. Gawad C, Koh W, Quake SR. Single-cell genome sequencing: current state of the science. Nat Rev Genet. 2016;17(3):175–88.

21. Bailey MH, Tokheim C, Porta-Pardo E, Sengupta S, Bertrand D, Weerasinghe A, et al. Comprehensive Characterization of Cancer Driver Genes and Mutations. Cell. 2018;173(2):371–85 e18.

22. DiRenzo J, Signoretti S, Nakamura N, Rivera-Gonzalez R, Sellers W, Loda M, et al. Growth factor requirements and basal phenotype of an immortalized mammary epithelial cell line. Cancer Res. 2002;62(1):89–98.

23. Jiang Y, Pjesivac-Grbovic J, Cantrell C, Freyer JP. A multiscale model for avascular tumor growth. Biophys J. 2005;89(6):3884–94.

24. Santoni D, Pedicini M, Castiglione F. Implementation of a regulatory gene network to simulate the TH1/2 differentiation in an agent-based model of hypersensitivity reactions. Bioinformatics. 2008;24(11):1374–80.

25. Wang Z, Birch CM, Sagotsky J, Deisboeck TS. Cross-scale, cross-pathway evaluation using an agent-based non-small cell lung cancer model. Bioinformatics. 2009;25(18):2389–96.

26. Davidich M, Bornholdt S. The transition from differential equations to Boolean networks: a case study in simplifying a regulatory network model. J Theor Biol. 2008;255(3):269–77.

27. Thomas R. Regulatory Networks Seen as Asynchronous Automata - a Logical Description. J Theor Biol. 1991;153(1):1–23.

28. Grieco L, Calzone L, Bernard-Pierrot I, Radvanyi F, Kahn-Perles B, Thieffry D. Integrative modelling of the influence of MAPK network on cancer cell fate decision. PLoS Comput Biol. 2013;9(10):e1003286.

29. Holt RC, Winter A, Schurr A, editors. GXL: Toward a standard exchange format. Reverse Engineering, 2000 Proceedings Seventh Working Conference on; 2000: IEEE.

30. Brandes U, Eiglsperger M, Herman I, Himsolt M, Marshall MS, editors. GraphML progress report structural layer proposal. International Symposium on Graph Drawing; 2001: Springer.

31. Wilensky U, Evanston I. NetLogo: Center for connected learning and computer-based modeling. Northwestern University, Evanston, IL. 1999:49–52.

32. Dent P. The multi-hit hypothesis in basal-like breast cancer. Cancer Biol Ther. 2013;14(9):778–9.

33. Pires MM, Hopkins BD, Saal LH, Parsons RE. Alterations of EGFR, p53 and PTEN that mimic changes found in basal-like breast cancer promote transformation of human mammary epithelial cells. Cancer Biol Ther. 2013;14(3):246–53.

34. Bessette DC, Tilch E, Seidens T, Quinn MC, Wiegmans AP, Shi W, et al. Using the MCF10A/MCF10CA1a Breast Cancer Progression Cell Line Model to Investigate the Effect of Active, Mutant Forms of EGFR in Breast Cancer Development and Treatment Using Gefitinib. PLoS One. 2015;10(5):e0125232.

35. Rivlin N, Brosh R, Oren M, Rotter V. Mutations in the p53 Tumor Suppressor Gene: Important Milestones at the Various Steps of Tumorigenesis. Genes & cancer. 2011;2(4):466–74.

36. Kastenhuber ER, Lowe SW. Putting p53 in Context. Cell. 2017;170(6):1062–78.

37. Eales KL, Hollinshead KE, Tennant DA. Hypoxia and metabolic adaptation of cancer cells. Oncogenesis. 2016;5:e190.

38. Wilson WR, Hay MP. Targeting hypoxia in cancer therapy. Nat Rev Cancer. 2011;11(6):393–410.

39. Harris AL. Hypoxia--a key regulatory factor in tumour growth. Nat Rev Cancer. 2002;2(1):38–47.

40. Haider S, McIntyre A, van Stiphout RG, Winchester LM, Wigfield S, Harris AL, et al. Genomic alterations underlie a pan-cancer metabolic shift associated with tumour hypoxia. Genome biology. 2016;17(1):140.

41. Grimes DR, Kelly C, Bloch K, Partridge M. A method for estimating the oxygen consumption rate in multicellular tumour spheroids. J R Soc Interface. 2014;11(92):20131124.

42. Macal CM, North MJ. Agent-Based Modeling and Simulation. Wint Simul C Proc. 2009:86-+.

43. Matthews RB, Gilbert NG, Roach A, Polhill JG, Gotts NM. Agent-based land-use models: a review of applications. Landscape Ecol. 2007;22(10):1447–59.

44. Wang Z, Butner JD, Kerketta R, Cristini V, Deisboeck TS. Simulating cancer growth with multiscale agent-based modeling. Semin Cancer Biol. 2015;30:70–8.

45. Thorne BC, Bailey AM, Peirce SM. Combining experiments with multi-cell agent-based modeling to study biological tissue patterning. Brief Bioinform. 2007;8(4):245–57.

46. Zanoni M, Piccinini F, Arienti C, Zamagni A, Santi S, Polico R, et al. 3D tumor spheroid models for in vitro therapeutic screening: a systematic approach to enhance the biological relevance of data obtained. Sci Rep. 2016;6:19103.

47. Chen CS, Mrksich M, Huang S, Whitesides GM, Ingber DE. Geometric control of cell life and death. Science. 1997;276(5317):1425–8.

48. Haeger A, Krause M, Wolf K, Friedl P. Cell jamming: collective invasion of mesenchymal tumor cells imposed by tissue confinement. Biochim Biophys Acta. 2014;1840(8):2386–95.

49. Ahmadzadeh H, Webster MR, Behera R, Jimenez Valencia AM, Wirtz D, Weeraratna AT, et al. Modeling the two-way feedback between contractility and matrix realignment reveals a nonlinear mode of cancer cell invasion. Proc Natl Acad Sci U S A. 2017;114(9):E1617–E26.

50. Garay T, Juhasz E, Molnar E, Eisenbauer M, Czirok A, Dekan B, et al. Cell migration or cytokinesis and proliferation?--revisiting the “go or grow” hypothesis in cancer cells in vitro. Exp Cell Res. 2013;319(20):3094–103.

51. Jimenez Valencia AM, Wu PH, Yogurtcu ON, Rao P, DiGiacomo J, Godet I, et al. Collective cancer cell invasion induced by coordinated contractile stresses. Oncotarget. 2015;6(41):43438–51.

